# Quantification of lysogeny caused by phage coinfections in microbial communities from biophysical principles

**DOI:** 10.1101/2020.04.22.056689

**Authors:** Antoni Luque, Cynthia Silveira

## Abstract

Temperate phages can integrate in their bacterial host genome to form a lysogen, often modifying the phenotype of the host. Lysogens are dominant in the microbial-dense environment of the mammalian-gut. This observation contrasts with the long-standing hypothesis of lysogeny being favored in microbial communities with low densities. Here we hypothesized that phage coinfections—the most studied molecular mechanism of lysogeny in lambda phage—increases at high microbial abundances. To test this hypothesis, we developed a biophysical model of coinfection and stochastically sampled ranges of phage and bacterial concentrations, adsorption rates, lysogenic commitment times, and community diversity from marine and gut microbiomes. Based on lambda experiments, a Poisson process assessed the probability of lysogeny via coinfection in these ecosystems. In 90% of the sampled marine ecosystems, lysogeny stayed below 10% of the bacterial community. In contrast, 25% of the sampled gut communities stayed above 25% of lysogeny, representing an estimated nine trillion lysogens formed via phage coinfection in the human gut every day. The prevalence of lysogeny in the gut was a consequence of the higher densities and faster adsorption rates. In marine communities, which were characterized by lower densities and phage adsorption rates, lysogeny via coinfection was still possible for communities with long lysogenic commitments times. Our study suggests that physical mechanisms can favor coinfection and cause lysogeny at poor growth conditions (long commitment times) and in rich environments (high densities and adsorption rates).

**Importance:** Phage integration in bacterial genomes manipulate microbial dynamics from the oceans to the human gut. This phage-bacteria interaction, called lysogeny, is well-studied in laboratory models, but its environmental drivers remain unclear. Here we quantified the frequency of lysogeny via phage coinfection—the most studied mechanism of lysogeny—by developing a biophysical model that incorporated a meta-analysis of the properties of marine and gut microbiomes. Lysogeny was found to be more frequent in high-productive environments like the gut, due to higher phage and bacterial densities and faster phage adsorption rates. At low cell densities, lysogeny via coinfection was possible for hosts with long duplication times. Our research bridges the molecular understanding of lysogeny with the ecology of complex microbial communities.

## Introduction

Temperate phages are viruses of bacteria that, upon infection, can integrate in their host’s genome as a prophage. This forms a symbiont with the bacterial host called lysogen. Half of bacterial genomes that have been sequenced contain prophages (Casjens, 2003; Fouts, 2006; Touchon *et al.*, 2016). Most lysogens display enhanced capabilities such as protection against other phage infections and additional metabolic pathways (Bondy-Denomy *et al.*, 2016; Jerlström Hultqvist *et al.*, 2018; Mavrich and Hatfull, 2019). Therefore, lysogeny has profound impacts in the structure and functioning of microbial communities. However, the drivers of lysogeny in microbial communities remain unclear.

The distribution of integrases, excisionases, and lambda-like repressor genes currently represents the best proxy to estimate the relative frequency of lysogeny in microbial communities (Luo *et al.*, 2020; Silveira *et al.*, 2020). The application of this approach to metagenomic analysis indicates that microbial-dense environments, such as the mammalian gut, are dominated by temperate phages and bacterial lysogens, compared with low-density marine ecosystems (Reyes *et al.*, 2010; Minot *et al.*, 2011, 2013; Beller and Matthijnssens, 2019; Shkoporov and Hill, 2019; Mirzaei *et al.*, 2020). In complete bacterial genomes from isolates, higher frequency of prophages is also observed in the gut (Anthenelli *et al.*, 2020). The gut-associated temperate phages mediate bacterial colonization dynamics in mouse models (Reyes *et al.*, 2013; Frazão *et al.*, 2019), and their distribution patterns have been associated with the presence of severe acute malnutrition (Reyes *et al.*, 2015).

Compared to animal-associated microbial communities, marine ecosystems have much weaker lysogenic signatures (Knowles *et al.*, 2016; Luo *et al.*, 2020). Genomic analyses have indicated that lysogeny is frequent in deep oligotrophic waters where bacterial abundances are low, ranging from 10^4^ to 10^6^ cells per ml (Muck *et al.*, 2014; Luo *et al.*, 2017). The increase in lysogeny in marine ecosystems with low productivity has been historically hypothesized to serve a low-density refugium for phages during poor host growth (Paul, 2008). Yet, high intrinsic growth rate is the most important predictor of the frequency of prophages in gut and marine bacteria with complete genomes sequenced (Lauro *et al.*, 2009; Touchon *et al.*, 2016). Lysogeny also increases in shallow marine environments where abundances of bacteria rise above 10^6^ cells/ml (Knowles *et al.*, 2016). The observations of lysogeny at both high and low productivity conditions suggest that an interplay between host physiology and density-dependence mechanisms modulate lysogeny in complex communities (Thingstad, 2000; Weitz *et al.*, 2015; Knowles *et al.*, 2016).

Phage coinfections are the best understood mechanism for the establishment of lysogeny in model phage and bacteria (Golding, 2016; Jimmy T. Trinh *et al.*, 2017). The percentage of lysogenized cells increase with the multiplicity of infection (MOI) in lambda phage and *E. coli* (Boyd, 1951; Lieb, 1953; Fry, 1959; Kourilsky, 1973; Herskowitz and Hagen, 1980). Coinfections increase the expression of the cII repressor lytic pathway (phage replication) and activate a cascade of genes responsible for phage integration (Kourilsky, 1975; Ptashne, 1986; Cheng *et al.*, 1988). Yet, the extent to which coinfections occur in complex microbial communities, and their impact on lysogeny has not been quantified.

After its first introduction, the term MOI became known in the field as the initial ratio of phage particles (P_0_) to bacterial cells (B_0_) added to an experiment (MOI = P_0_/B_0_). This definition of MOI, however, does not capture the effective number of coinfections, which also depends on the chances of encounter between the phage and the host (Joiner *et al.*, 2019). A recent stochastic model shows that at the single-cell level, the average number of coinfections is primarily determined by the phage concentrations and phage decision times (Lang *et al.*, 2020). Since high productive environments have higher phage concentrations, here we hypothesize that the prevalence of lysogeny in these environments is a consequence of an increase in coinfections.

To test this hypothesis, a biophysical model was derived to incorporate the physical traits that determine phage coinfections (COI) and its associated probability of lysogeny. The model was simulated for a range of phage and bacterial abundances, community diversity, adsorption rates, and lysogenic commitments times from marine and mammalian gut communities. The availability of large amounts of public data from these two ecosystems allowed to test the hypothesis across a wide range of microbial densities.

## Methods

### Physical model of the probability of lysogeny through phage coinfection

The average number of phage coinfections (COI) was derived from physical properties of phage and bacteria (Figure 1a). The rate of phage infections on a single bacterium can be estimated by solving the Smoluchowski coagulation equation (Joiner *et al.*, 2019). In steady state, this rate is the product of the phage concentration (P) and the phage adsorption rate (*α*), which depends on the mobilities and sizes of both the phage particle and the bacterium. The number of coinfections is defined by the number of phages infecting a cell within a given time window. This number is the product of the infection rate and the time window. In the case of lysogeny, this time corresponds to the lysogenic commitment time (*τ*) (Jimmy T. Trinh *et al.*, 2017). This led to the average phage coinfection equation (COI)

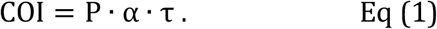

**Figure 1.**
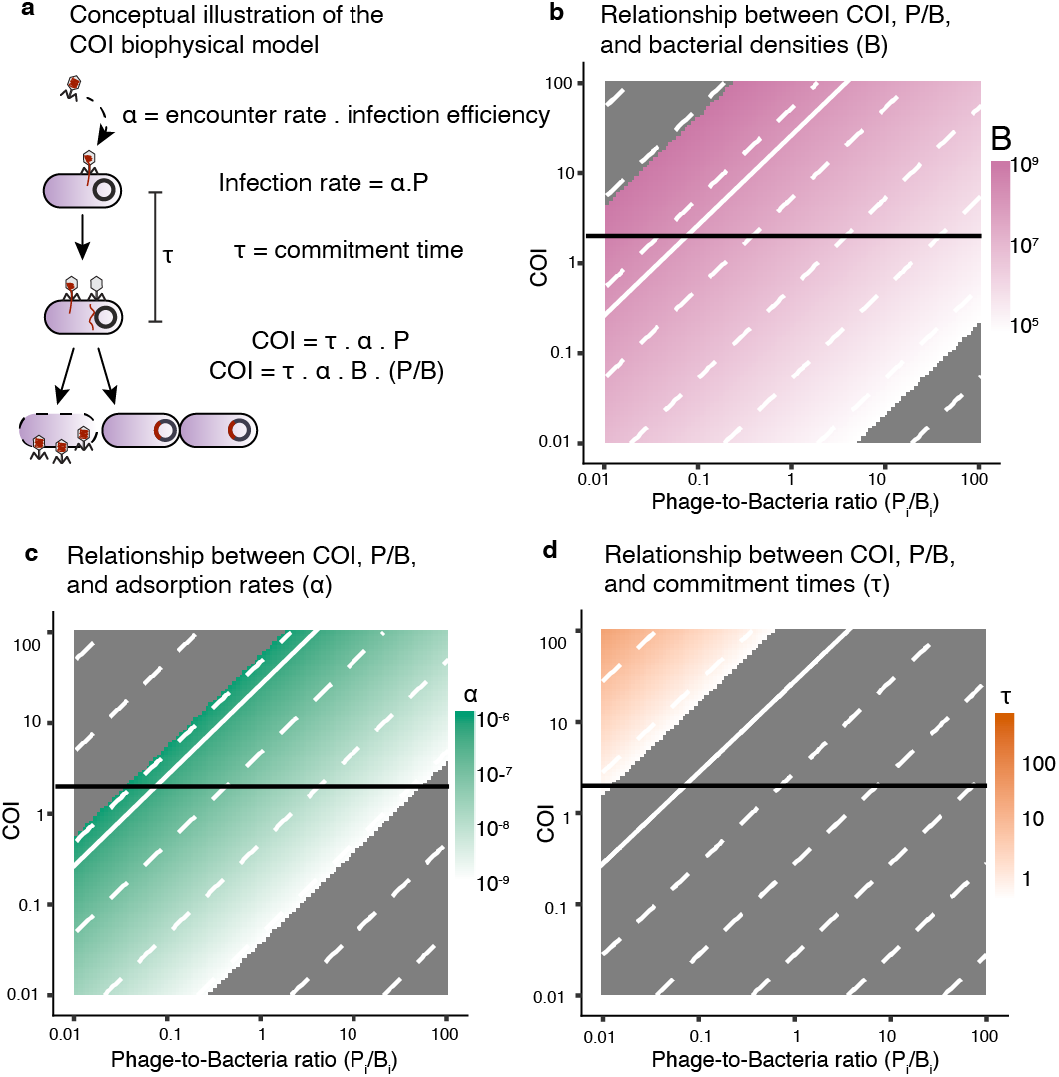
Relationship between number of coinfections and phage-to-bacteria ratios. **a**) Illustration of the derivation and parameter description for the biophysical coinfection (COI) model, Eq. (1). Relationship between COI and phage-to-bacteria ratios as a function of bacterial abundances (**b**), adsorption rates (**c**), and lysogenic commitment times (**d**). The color gradients cover the environmental ranges of values for each of these parameters: bacterial concentrations (pink), adsorption rates (green), and lysogenic commitment times (orange). (**b-d**) The dotted white lines indicate constant values of the parameters in the gradient scale. The grey areas correspond to values beyond the environmental ranges of bacterial concentrations, adsorption rates, and commitment times obtained from the meta-analysis of marine and gut ecosystems. The horizontal black line indicates COI = 2. The solid white line indicates the median values for the bacterial concentration (B_0_), adsorption rates (*a*_0_), and lysogenic commitment time (*τ*_0_) for lambda.

The average coinfection (COI) was also expressed as a function of the phage-to-bacteria ratio, COI = α · τ · B ∙ (P/B), for one single phage-host pair to facilitate the comparison with lambda MOI experiments. Here, B was the bacterial concentration. COI was studied numerically as a function of the phage-to-bacteria ratio (P/B) for the range 0.01-100. The median values extracted for the lambda adsorption rate (*α*_0_ = 5.6 10^−7^ ml/h), lysogenic commitment time (*τ*_0_ = 0.1 h), and bacterial centration (*B*_0_ = 5 10^8^ cells/ml) were used as reference values (see Methods section on meta-analysis for lambda parameters). Two parameters were fixed at these reference values, and the third was explored over a range of values. These ranges were 10^5^ to 10^10^ cells/ml for bacterial concentrations, 10^−11^ to 10^−6^ ml/h for the phage adsorption rates, and 10^−3^ to 10^2^ h for the lysogenic commitment times.

A lysogen was formed when a cell was infected within the lysogenic commitment time by two or more phages from the same pair. This was based on the effect of cooperative infection by lambda phages on the production of lysogens (Zeng *et al.*, 2010; Golding, 2016; Jimmy T. Trinh *et al.*, 2017). Thus, the probability of lysogeny, p_1ys_, was determined by p_1ys_= 1 − p(0) − p(1), where p(k) was the probability of having k coinfections within the commitment time. The probability of coinfections was calculated by assuming that each infection was independent, and that the phage concentration (P), phage adsorption rates (*α*), and lysogenic commitment times (*τ*) remained constant. Following these assumptions, the probability of k coinfections was given by a Poisson process with average coinfection COI, Eq. (1), leading to the equation p(k) = COI^k^e^−COI^/k!. Thus, the probability of forming a lysogen via coinfection was

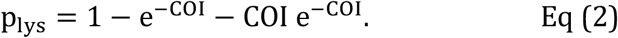

This probability of lysogeny was compared with the percentage of lysogeny obtained from lambda and *E. coli* MOI experiments (Fry, 1959; Kourilsky, 1973). The empirical data for percentage of lysogeny and MOI was fitted using non-linear least squares method for Hill-Langmuir cooperation models, *f*(*x*) = *ax*/(*b* + *x^n^*), with cooperation orders n=1, n=2, and n=3.

### Meta-analysis of marine and gut microbiomes

#### Adsorption rates

The adsorption rates were obtained from prior experiments for 71 phage-host pairs from marine and gut microbiomes (Source data 1). The data consisted of values for tailed phages infecting *E. coli* (Schwartz, 1976; Joiner *et al.*, 2019), *Synechococcus sp.* (Waterbury and Valois, 1993; Mann, 2003; Stoddard *et al.*, 2007), *Prochlorococcus sp.* (Avrani *et al.*, 2011; Schwartz and Lindell, 2017; Beckett *et al.*, 2019), *Vibrio sp.* (Johnson, 1968; Levisohn *et al.*, 1987; Cohen *et al.*, 2013; Traits *et al.*, 2017), *Roseobacter sp.* (Huang *et al.*, 2010), and *Pseudoalteromonas sp.* (Deng *et al.*, 2013). A t-test (double tailed) compared the marine and gut values.

#### Lysogenic commitment times

The lysogenic commitment time (*τ*) was assumed to be 20% of the bacterial duplication time (Cortes *et al.*, 2017; Jimmy T. Trinh *et al.*, 2017). The ranges of bacterial duplication times at the community level were obtained *in situ* for marine ecosystems (Kirchman, 2016) and *in vivo* for mammalian gut ecosystems (Heinken *et al.*, 2013; Myhrvold *et al.*, 2015; Vandeputte *et al.*, 2016) (Source data 2).

#### Viral-like particles and cell abundances

Direct counts of viral-like particles (VLPs) and microbial cells were obtained for marine surface waters (Parsons *et al.*, 2012; Knowles *et al.*, 2016) and animal-associated microbiomes (Hoyles *et al.*, 2008; Lepage *et al.*, 2008; Furlan, 2009; Kim *et al.*, 2011; Barr *et al.*, 2013) (Source data 3). A t-test (double tailed) was applied to compare the concentrations of VLPs and cells between marine and animal ecosystems.

#### Phage and bacterial diversities

The rank-abundance curves of the frequency of phage genotypes representing unique viral contigs at 98% sequence identity were reconstructed from the median slope and intercept of power-law functions fitted to 192 marine viromes and 1,158 human-associated viromes (Cobián Güemes *et al.*, 2016) (Source data 4). The rank-abundance curves of the frequency of bacterial species in marine communities were obtained from operational taxonomic unit (OTU) tables constructed by clustering universal, protein-coding, single-copy phylogenetic marker genes into metagenomic OTUs (which can be interpreted as species-level clusters) from the Tara Oceans dataset (Source data 5) (Mende *et al.*, 2013; Sunagawa *et al.*, 2013). For animal-associated bacterial microbiomes, rank-abundance curves were constructed using OTU tables obtained by mapping metagenomic reads from 11,850 human gut-associated metagenomes to 92,143 metagenome-assembled genomes (Almeida *et al.*, 2019). Consensus rank abundance curves were obtained by averaging the frequency of bacteria in the same rank across the metagenomes of marine and gut associated microbiomes, respectively.

### Quantification of lysogeny through phage coinfection in communities

The biophysical COI model, Eqs. (1) and (2), was applied to predict the probability of lysogeny in marine and gut ecosystems as a result of coinfections. The model generated stochastic communities that sampled empirical phage and bacteria concentrations, relative abundance of the top 100 members of the community, phage adsorption rates, and lysogenic commitment times obtained from the meta-analysis of marine and gut ecosystems described above.

#### Stochastic sampling

The model generated 100,000 stochastic communities for both marine and gut ecosystems using latin hypercube sampling (LHS). For each ecosystem, the range of bacterial concentrations, adsorption rates, and lysogenic commitment times were divided in 100,000 equidistributed values in logarithmic scale (base 10). 100,000 random values were selected from each range and combined without repeating any value, that is, all equidistributed values were sampled, following the standard LHS implementation (McKay *et al.*, 1979; Weitz *et al.*, 2017; Anthenelli *et al.*, 2020)

#### Parameter ranges

The ranges of bacterial concentrations used were 3.78·10^4^ – 6.75·10^6^ bacteria/ml (marine) and 3.45·10^5^ – 7.60·10^9^ bacteria/ml (gut). The ranges of phage concentrations used were 1.45·10^5^ – 3.80·10^7^ phages/ml (marine) and 5.09·10^6^ – 1.05·10^10^phages/ml (gut). The ranges of phage adsorption rates used were 7.2 10^−10^ – 3.7 10^−7^ ml/h for marine and 5.9 10^−8^ – 1.2 10^−6^ for gut. The ranges of lysogenic commitment times used were 11 h – 808 h for marine and 2.74 h –7.27 h for gut.

#### Assumed relationships

The phage concentration (P) was assumed to follow a power function relationship with the bacterial concentration (B)(Knowles *et al.*, 2016; Anthenelli *et al.*, 2020): P(B) = a (B/B_u_)^b^, where the bacterial concentration was given in units of B_u_ = bacteria/ml. The prefactor a and exponent b were obtained by fitting the power function to the viral and microbial counts obtained in the marine and gut meta-analysis. A linear regression fit was applied using least-squares method to the log-log data in base 10. The parameters obtained were a = 10^2.50^ and b = 0.712 for marine and a = 10^5.35^ and b = 0.388 for gut. To reproduce the noise observed in empirical viral communities, the value P(B) was weighted by a normal distribution, N(mean, S.D.), with mean 1 and standard deviation 0.05, that is, P = P(B)*N(1,0.05). The final value of the phage concentration was constrained within the empirical phage range, that is, P_min_ ≤ P ≤ P_max_. The community model also assumed that the most dominant phages infected the most dominant bacteria (Minot *et al.*,.2011, 2013; Coutinho *et al.*, 2017). This led to a phage-host network where the phage of rank *i* infected the bacteria with the same rank *i*.

## Results

### Relationship between COI and phage-to-bacteria ratios

The model introduced here predicted the average number of phage coinfections (COI) occurring across a gradient of phage-to-bacteria ratios for single pairs (Figure 1a). To illustrate the relationship of COI with different physical parameters, the typical laboratory values for lambda were used as a reference, and only one value was varied at a time covering typical environmental ranges (see meta-analysis section for environmental ranges). Coinfection (COI) increased with phage-to-bacteria ratio (P_i_/B_i_) (Figure 1). The level of COI was higher for higher bacterial concentration (Figure 1b), phage adsorption rate constant (Figure 1c), and lysogenic commitment time (Figure 1d). The model showed that an average of two or more lambda coinfections (COI ⩾ 2) were unlikely to occur at bacterial densities below 10^6^ cells/ml, even for phage-to-bacteria ratios above 10 (bottom right values in Figure 1b). The typical adsorption rates for lambda were near the upper bound of coinfections generated per phage-to-bacteria ratio (Figure 1c). Due to the short lysogenic commitment times of lambda, the average coinfections per phage-to-bacteria ratio unit were below the environmental values (Figure 1d).

The probability of lysogeny as a function of average coinfections (COI) was compared with lambda-*E. coli* MOI experiments, where MOI was defined operationally as the initial phage-to-bacteria ratio. The percentage of lysogenized cells increased as a function of MOI and was best described by a sigmoidal Hill-Langmuir equation of order n = 2, compared to orders n = 1 and n = 3 (Figure 2a). This equation implied that two phages cooperated in producing lysogeny, in agreement with single-molecule experiments (Jimmy T Trinh *et al.*, 2017). This empirical model was functionally similar to the predicted probability of lysogeny from the average coinfection Poisson model, Eq. (2), which assumed that at least two co-infections were necessary to produce lysogeny (Figure 2b). The discrepancy between the maximum percentage of lysogeny for the MOI and COI models was due to the fact that the MOI model was a function of the initial phage-to-bacteria ratio instead of the effective number of coinfections per cell.

**Figure 2.**
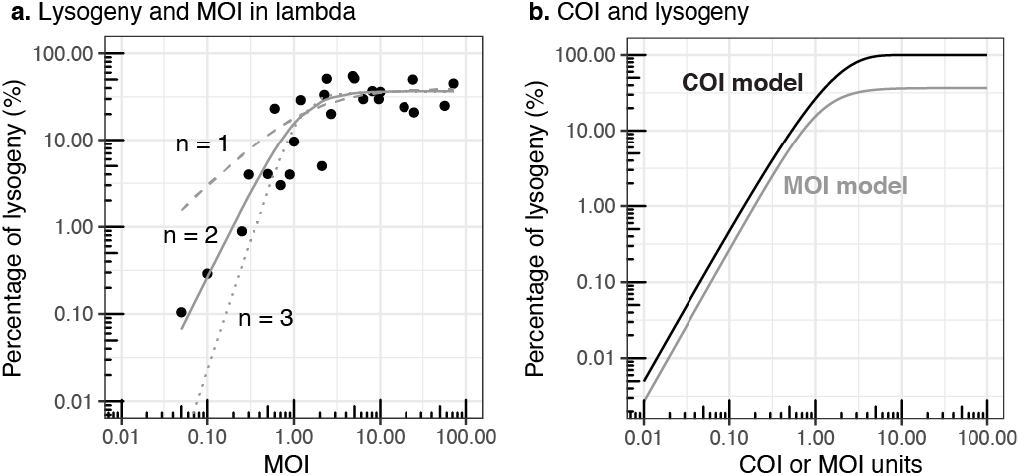
Comparison of percentage of lysogeny for lambda MOI and COI model. **a)** Relationship between the percentage of lysogeny and MOI (initial phage-to-bacteria ratio) in lambda-E *. coli* experiments (Fry, 1959; Kourilsky, 1973). The lines correspond to fitted Hill-Langmuir cooperation models of order n =1 (dashed), n=2 (solid), and n=3 (dotted). **b)** Percentage of lysogeny estimated from the coinfection probability model as a function of COI, Eq. (2) (solid black line). Hill-Langmuir model of order n=2 from panel **a**) as a function of MOI (grey solid line).

### Meta-analysis of COI physical parameters from marine and animal ecosystems

To apply the biophysical COI model to microbial communities and estimate lysogeny generated by phage coinfections, the ranges of phage adsorption rates, lysogenic commitment times, and phage-bacteria pair abundances were determined for marine and animal ecosystems. The range of adsorption rates was 7.2 10^−10^ – 3.7 10^−7^ ml/h for marine phages infecting *Prochlorococcus, Roseobacter, Pseudoalteromonas, Synechococcus, and Vibrio* (Figure 3a). The range of adsorption rates was 5.9 10^−8^ – 1.2 10^−6^ ml/h for gut phages infecting *E. coli*. The median adsorption rates for gut phages was one order of magnitude higher (median 4.21 10^−7^ ml/h) than the median for the marine phages (median 3.43 10^−8^ ml/h, Figure 3c, t-test p = 7.23 10^−10^). The ranges of estimated lysogenic commitment times were much longer in marine communities, 11-808 h, than in mammalian gut, 2.74–7.27 h. This was a consequence of the long duplication times of marine communities in their natural environment (Source data 5). The phage and bacteria pair abundances were determined by combining the total and relative abundances of phage and bacteria in each ecosystem. The total abundances were at least two orders of magnitude higher in animal-associated mucosa compared to the free-living communities of surface marine environments (t-test p = 7.02 10^−15^ for phage, Figure 3b, and p-value = 4.17 10^−7^ for bacteria, Figure 3c). For phages, the ranges were 1.4 10^5^–3.7 10^7^ phages/ml (marine) and 5.1 10^6^–1.1 10^10^ phages/ml (animal), while bacteria ranged from 3.8 10^4^ to 6.8 10^6^ cells/ml (marine) and from 3.5 10^5^ to 7.7 10^9^ cells/ml (animal) (Figures 3b and 3c). The most abundant phage genotype (P_1_) in marine environments comprised only 0.8% of the total phage community, while in the gut, the dominant phage comprised just over 1% of the community (Figure 3d). In the bacterial community, this pattern was inverted, with the dominant bacterial species (B_1_) reaching 19% in marine environments, but only 15% in the gut (Figure 3e).

**Figure 3.**
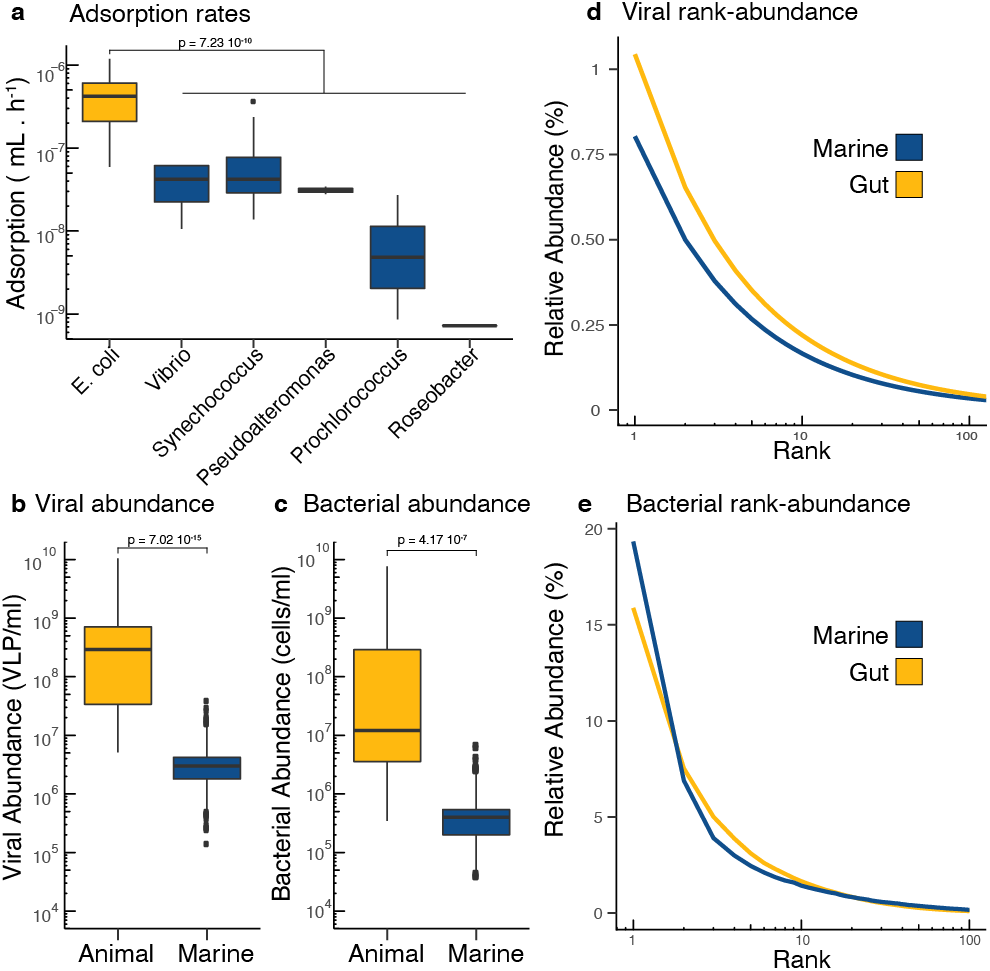
Meta-analysis of COI parameters in marine and animal microbiomes. **a)** Adsorption rates from single phage-host pairs of bacterial hosts derived from the mammalian gut (yellow) and marine environments (blue). The same color coding applies to all panels. **b)** Viral abundances and **c)** bacterial abundances per ml in animal-associated and marine environments. **d)** Rank abundance curve displaying the top 100 ranks of phage genotypes identified in metagenomic data from human gut and marine samples. **e)** Rank abundance curve displaying the top 100 ranks of bacterial genotypes identified in metagenomic data from human gut and marine samples. The references for each original study and the values for each datapoint are provided as Source Data 1 to 5.

### Estimated lysogeny by coinfection in microbial communities

The community model assumed a direct phage-host network, where each phage rank infected the same rank level in the bacterial community, that is, P_i_ infected B_i_ (Figure 4a). The biophysical COI model quantified the percentage of lysogeny generated by phage coinfections for each pair by stochastically sampling the parameter ranges from the meta-analysis of marine and gut ecosystems. The percentage of lysogeny caused by coinfections increased with total bacterial density (Figures 4b and S.1). Lysogeny was more frequent in the gut, where 25% of the simulated communities displayed at least 25% of bacteria becoming lysogens by coinfection (Figure 4c and Tables S.1 and S.2). Instead, 90% of the marine communities displayed 10% or less bacteria becoming lysogens by coinfection (Figure 4c and Tables S.1 and S.3). Among communities with a percentage of lysogeny larger than 1%, the most abundant phage-host pairs contributed an average of 67±12% (S.D.) for marine and 51±16% for gut to the total lysogeny (Figures 4d and S.2). This was significantly higher than the contribution from the second most abundant phage-host pair, which yielded 13±1% for marine and 15±2% for gut. For communities with lysogeny above 1%, the most abundant phage-host rank displayed median average coinfections of COI = 1.00 for marine (Figures 4e and S.2 and Table S.4) and COI = 2.35 for gut (Figure 4f and S.2 and Table S.5).

**Figure 4.**
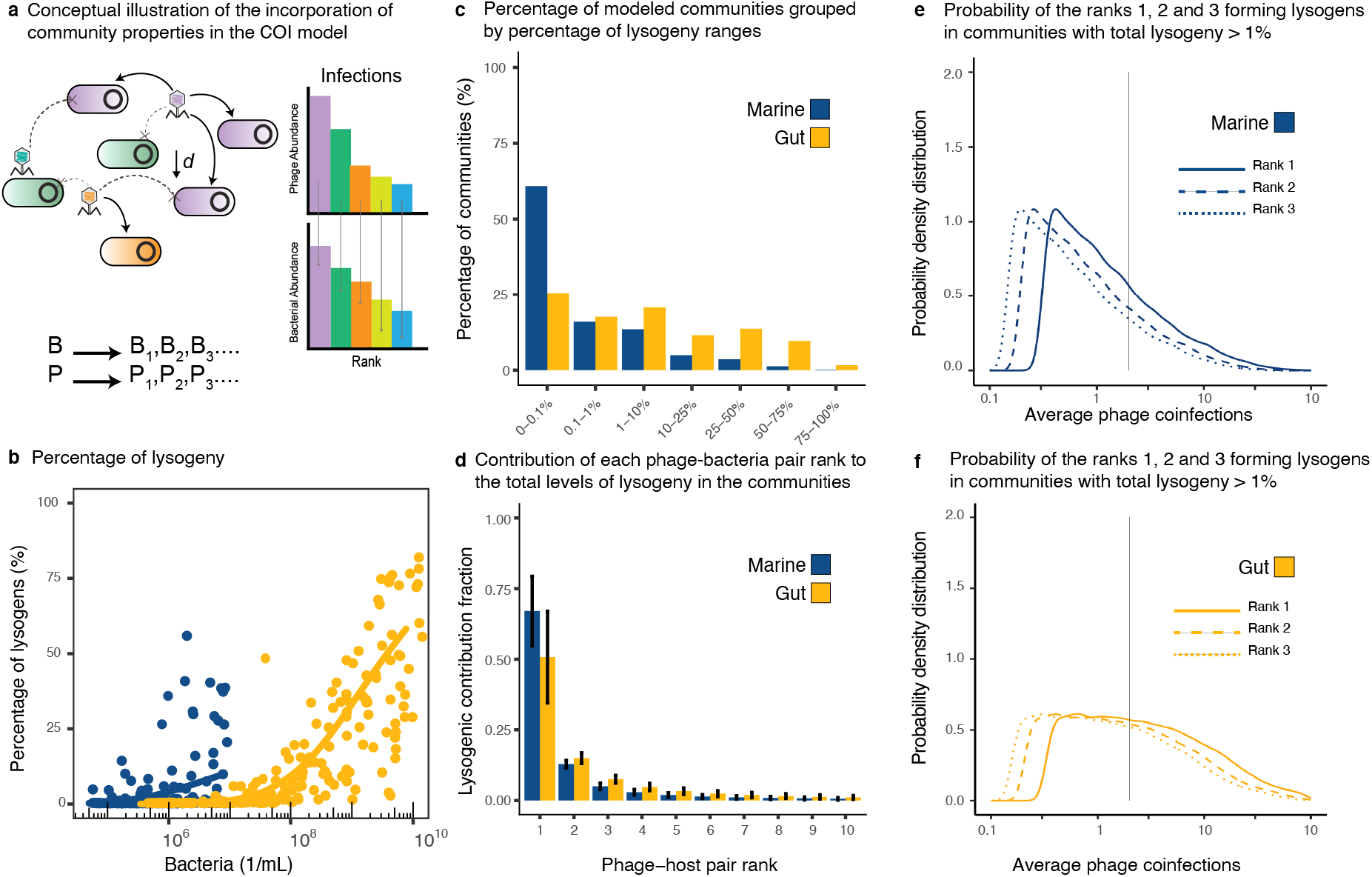
Lysogeny caused by coinfections in communities. **a)** Conceptual figure describing the implementation of the biophysical COI model with stochastic sampling of community parameters. Each Phage P_i_ infected each bacterium B_i_ according to their ranks in the community. The chances of encounter decreased with phage-host rank as a function of their absolute abundances in the community. **b)** Percentage of lysogenic cells in the bacterial community predicted to be formed through phage coinfections as a function of the total bacterial density. The data points are a subsample of 200 stochastic models out of 100,000 sampled models for each ecosystem. The solid lines represent generalized additive models (GAM) fitted to the full data set for marine (blue) and gut (yellow) ecosystems. **c)** ercentage of sampled communities displaying different ranges of lysogeny. **d)** Average contribution (with error bars corresponding to S.D.) from the top phage-bacteria pair ranks to the lysogenic pool. **e-f)** Probability distributions of average phage coinfections (COI) for the top ranks in communities with lysogeny above 1% for marine and gut ecosystems. The vertical line indicates COI = 2.

### Physical parameters contributing to the formation of lysogens in communities

Communities with at least 1% lysogeny caused by coinfection were analyzed to extract the distribution of physical parameters associated to lysogeny. The distribution of bacterial abundances favoring lysogeny in marine communities was skewed towards high densities with a median of 1.5 10^6^ cells/ml (Figure 5a and Table S.4). In gut communities, low bacterial abundances did not contribute to lysogeny, displaying a first quartile at 3.8 10^5^ cells/ml (Figure 5a and Table S.5). Phage concentrations in marine communities were also skewed towards higher densities (median of 9.1 10^6^ phages/ml) but more centered compared to the bacterial density distribution (Figure 5b and Table S.4). In the gut, low phage concentrations did not contribute to lysogeny, displaying a first quartile at 2.8 10^8^ phages/ml (Figure 5b and Table S.5).

**Figure 5.**
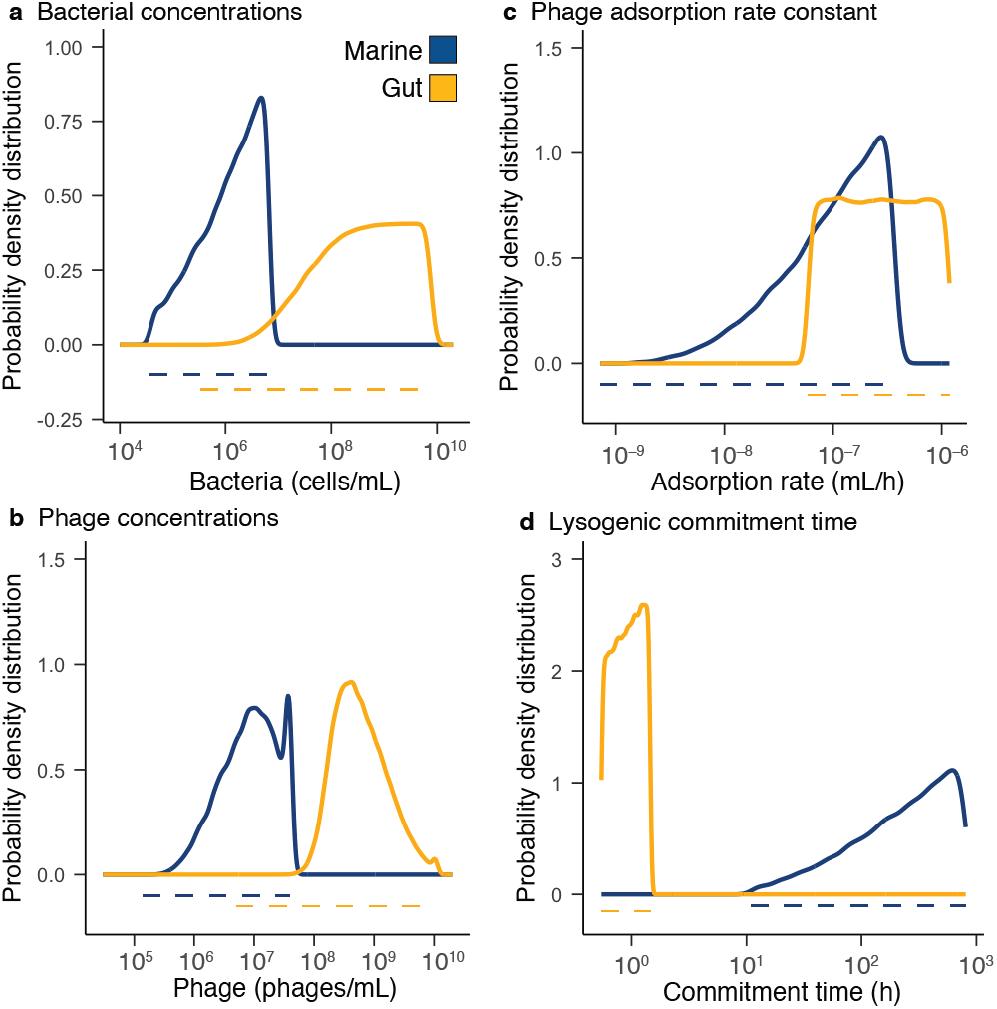
Ranges of parameter values leading to lysogeny caused by coinfection. Probability density distributions (solid lines) for bacterial abundances **(a)**, phage densities **(b)**, adsorption rates **(c)**, and lysogenic commitment times **(d)** in marine (blue) and gut (yellow) communities displaying lysogeny levels above 1%. The dotted lines indicate the range of parameters explored in the model and obtained from the meta-analysis of each ecosystem.

Phage adsorption rates in marine communities producing lysogeny were skewed towards high values with a median of 1.1 10^−7^ ml/h (Figure 5c and Table S.4). In the gut, instead, the full range of adsorption rates contributed to communities with lysogeny (Figure 5c and Table S.5). The lysogenic commitment time in marine communities producing lysogeny was again skewed towards long window times, with a median of 262 h (Figure 5d and Table S.4). For the gut, the full range of lysogenic commitment times contributed to communities with lysogeny, but larger values had higher likelihood of contribution, displaying a median lysogenic commitment time of 0.92 h (Figure 5d and Table S.5).

## Discussion

The stochastic biophysical COI model estimated an increase of coinfections and percentage of lysogeny in high-productive mammalian gut microbial communities (Figure 4). This higher frequency of coinfections explained the dominance of lysogenic bacteria observed in animal mucosa microbiomes (Reyes *et al.*, 2010; Minot *et al.*, 2013; Silveira and Rohwer, 2016; Shkoporov and Hill, 2019; Anthenelli *et al.*, 2020; Mirzaei *et al.*, 2020). The model also indicated that the formation of lysogens through coinfection was compatible with the observation of low virus-to-microbe ratios (VMR < 1) in high-density animal mucosa (Kim *et al.*, 2011; Mirzaei *et al.*, 2020) (Figures 3 and 5). This was possible because coinfections were not required to occur simultaneously. Instead, they occurred within the lysogenic commitment time, which was assumed to be proportional to the duplication time of bacteria *in vivo* (Heinken *et al.*, 2013; Myhrvold *et al.*, 2015; Vandeputte *et al.*, 2016).

The lysogenic commitment time played a paramount role in the findings of the model because the duplication time of gut bacteria *in vivo* is significantly slower than the duplication time of gut bacterial isolates in pure cultures (Heinken *et al.*, 2013; Myhrvold *et al.*, 2015; Vandeputte *et al.*, 2016). For laboratory *E. coli* duplication times, the model predicted average number of coinfections almost an order of magnitude lower than *in vivo* (Figure 1d). This result aligned with the low number of coinfections observed even at higher MOIs in a recent agent-based stochastic model that used similar parameters to lambda and *E. coli* (Lang *et al.*, 2020). The influence of the lysogenic commitment time was even more pronounced in marine communities (Figure 5d). Levels of lysogeny above 10% were only possible for a small fraction of communities that displayed long lysogenic commitment times, which compensated the lower adsorption rates and phage and bacteria abundances in marine communities (Figure 3 and 5).

The relationship between the lysogenic commitment time and duplication time in the biophysical COI model was based on lambda-*E. coli* single-cell experiments, which identified the relationship *τ* ~ 20% of the duplication time (Cortes *et al.*, 2017; Jimmy T. Trinh *et al.*, 2017). This pattern arises from the voting phenomenon, where each coinfecting phage genome independently votes for lytic or lysogenic commitment (Zeng *et al.*, 2010). In cells that undergo lysogeny phages cooperate for host cell resources (Jimmy T. Trinh *et al.*, 2017). Late phage genome arrivals contribute less to the cell-level decision, and the time window for the contribution of the second phage is proportional to the transcription level of cII repressor, which varies with the host growth rate (Cortes *et al.*, 2017). Future studies addressing the relationship between the lysogenic commitment time and host duplication time in other phages would facilitate the refinement of the model and further test its validity in different ecosystems.

The stochastic community simulations assumed that the most dominant phages infected the most dominant bacteria (Minot *et al.*, 2011, 2013; Coutinho *et al.*, 2017). This assumption allowed to bridge the environmental data on total viral and bacterial abundances with the species distributions from genomic data (Figure 3). It resulted into a higher contribution from the dominant ranks to lysogeny (Figure 4d). While the positive relationship between phage and host abundance has been shown for some dominant species in marine and gut samples (Deng *et al.*, 2013; Zhao *et al.*, 2013; Džunková *et al.*, 2019), the prediction of phage hosts from genomic data is a current challenge, and the host of most phages identified through metagenomics remain unknown (Edwards *et al.*, 2016; Gregory *et al.*, 2019; Silveira *et al.*, 2020). The reconstruction of accurate phage-bacterial infection networks will in the future improve the models’ predictive power (Labonté *et al.*, 2015; Marbouty *et al.*, 2017).

Only 10% of marine communities simulated displayed levels of lysogeny above 10%. These communities were characterized by high phage and bacterial concentrations for marine microbiomes, displaying medians of 1.3 10^7^ phages/ml and 2.0 10^6^ cells/ml (Figures 5a and 5b). This finding was consistent with the weak signals of lysogeny observed in marine ecosystems (Knowles *et al.*, 2016; Luo *et al.*, 2020). The average percentage of lysogens formed by coinfection in the community decreased by almost two orders of magnitude as total bacterial concentrations reduced from 10^6^ to 10^5^ cells/ml (Figure S.1b). Phage coinfection, thus, could not explain an increase of lysogeny at low cell densities. Viral metagenomic studies from deep oceans with microbial abundances ranging from 10^4^ to 10^5^cells per ml showed an increase in the genomic evidence for lysogeny compared to more productive surface waters (Coutinho *et al.*, 2019; Luo *et al.*, 2020). In these environments, lysogeny has been proposed to serve as low-density refugium for temperate phages in conditions of poor host growth and scarce resources for viral particle production (Jiang and Paul, 1994; Maurice *et al.*, 2010; Paul and Weinbauer, 2010; Brum *et al.*, 2015). In the biophysical COI results, lysogeny in these low cell density communities was only possible at very long commitment times. Additional mechanisms might be necessary to support the increase of lysogeny at these low concentrations. One explanation could be the favored phage integration in starved cells as observed in lambda due to the reduced degradation of the cII repressor (Shotland *et al.*, 1997).

Metabolic shifts might also be reinforcing the formation of lysogens at high cell abundances. High energy availability modifies the bacterial central carbon metabolism by decoupling growth rates from growth efficiencies (Russell and Cook, 1995; Haas *et al.*, 2016; Silveira *et al.*, 2019). The metabolic decoupling, which is common is hypoxic and anaerobic environments, decreases the ATP yield turning the intracellular environment more favorable for lysogeny. As a result, high-productive environments may favor lysogeny by both phage coinfections and the decrease in ATP yield. The incorporation of metabolic mechanisms was beyond the scope of the current biophysical model but will improve predictions of lysogeny in the future.

## Conclusion

The stochastic biophysical COI model proposed here identified the ranges of physical parameters that drive phage coinfections in complex microbial communities. The model predicted high frequency of lysogeny caused by phage coinfections in the high-density mammalian gut. This finding was a consequence of high phage and bacterial densities and fast phage adsorption rates in comparison with marine communities. Longer lysogenic commitment times *in vivo* when compared to laboratory isolates also contributed to high-lysogeny in the gut. The simulated marine communities showed lower frequency of lysogeny by coinfections. Those communities that displayed a high fraction of lysogeny were characterized by long lysogenic commitment times. Our findings bridge the main molecular mechanism causing lysogeny in laboratory systems with metagenomic observations of lysogeny in complex microbial communities.

